# Self-supervision deep learning models are better models of human high-level visual cortex: The roles of multi-modality and dataset training size

**DOI:** 10.1101/2025.01.09.632216

**Authors:** Idan Daniel Grosbard, Galit Yovel

## Abstract

With the rapid development of Artificial Neural Network based visual models, many studies have shown that these models show unprecedented potence in predicting neural responses to images in visual cortex. Lately, advances in computer vision have introduced self-supervised models, where a model is trained using supervision from natural properties of the training set. This has led to examination of their neural prediction performance, which revealed better prediction of self-supervised than supervised models for models trained with language supervision or with image-only supervision. In this work, we delve deeper into the models’ ability to explain neural representations of object categories. We compare models that differed in their training objectives to examine where they diverge in their ability to predict fMRI and MEG recordings while participants are presented with images of different object categories. Results from both fMRI and MEG show that self-supervision was advantageous in comparison to classification training. In addition, language supervision is a better predictor for later stages of visual perception, while image-only supervision shows a consistent advantage over a longer duration, beginning from 80ms after exposure. Examination of the effect of data size training revealed that large dataset did not necessarily improve neural predictions, in particular in visual self-supervised models. Finally, examination of the correspondence of the hierarchy of each model to visual cortex showed that image-only self-supervision led to better correspondence than image only models. We conclude that while self-supervision shows consistently better prediction of fMRI and MEG recordings, each type of supervision reveals a different property of neural activity, with language-supervision explaining later onsets, while image-only self-supervision explains long and very early latencies of the neural response, with the model hierarchy naturally sharing corresponding hierarchical structure as the brain.

## Intro

Visual processing in the human brain is a distributed process, performed by hierarchical processing of visual features, starting from “simple” features identified in small receptive fields [1,2] to more complex features spanning the entire visual field [3–6]. This neural process has been studied both spatially and temporally, where information propagates from the early visual cortex (EVC) at early stages of processing (∼80-120ms) to the ventral visual stream (VVS) in later stages (∼150-250ms) with increased complexity and specificity of response to visual domains [7,8].

In an attempt to decipher the information that is represented in the different spatio-temporal processing stages, recent studies turned to Artificial Neural Networks (ANN), where processing also takes place in a hierarchical manner [9–13]. Early findings focused on identifying a shared hierarchy of emergent features in both regions of cortex as identified in fMRI, as well as in temporal emergence as measured in MEG [9,10]. Beyond showing great promise for predicting neural activity, Cichy and Kaiser [14] had identified the possibilities of ANN based models in explaining neural activity in terms of teleological explanation, i.e. allowing us to explain neural response in terms of the goal they serve to perform cognitive functions.

Consequentially, in recent years much research has attempted to isolate the specific components in learning that facilitates ANN convergence to perform similarly to humans, with much focus on architecture, optimization objectives, and optimization rules [15]. For example, Storrs et al. [16] have examined the effects of model architectural differences on their ability to predict neural response to images, finding that specific choice in hypothesis class (e.g., model architectures) had little effect on the ability to predict neural response to images, as long as they were subtypes of convolutional neural network (CNN) trained on natural images. Ratan Murty et al. [17] showed that training on specific image domains such as faces or places will increase the ability of a model to predict domain specific neural regions, such as the fusiform face area (FFA) or the parahippocampal place area (PPA). Dwivedi et al. [18] further increased the differentiation by examining the differences between models trained on the same dataset, to excel at different objectives. Specifically, they identified that models trained on two-dimensional tasks such as autoencoding, colorization, inpainting, and attention to 2-dimensional details were most predictive of the EVC, models trained to predict semantic properties showed best performance in predicting the inferior-temporal (IT) cortex, and models trained on objectives reliant on 3-dimensional tasks such as reshading, creating depth maps and calculating distances were best predictive of neural activity in the dorsal stream. Jozwik et al. [19] showed that more anterior brain regions, and neural responses at later onsets, are better predicted using semantic models manually curated visual and semantic features rated by human participants, while posterior regions and earlier activations are better predicted by image-classification CNN-based models. Deveraux et al. [20] had shown similar findings where the semantic model was based on ANN trained on semantic attraction.

As artificial and biological neural networks are independent systems, these similarities in structure and representation do not pose as proof that the neural mechanisms stem from the same optimization as ANNs. However, due to the unprecedented ability to predict neural activity, philosophical frameworks were used to analyse what could be learned about the brain from models. One such framework is the Neuroconnectionist research program [21] proposing that, under some axioms of shared computational basis, by merely identifying success progression of ANN-neuro research that relies on specific model characteristics points towards that characteristic being shared by the brain. Furthermore, Simony et al [22] recently proposed a novel perspective for understanding these similarities with the perspective of convergent evolution. Using the perspective of common characteristics of visual models that perform well when predicting visual models are bound together and viewed as those that are critical to create potent visual systems, thus this race for identifying better predicting models can prove to be insightful for understanding the underlying neural mechanisms of perception.

The similarities between humans and ANNs have been primarily studied with ANNs that were optimized for object recognition using supervised learning. In supervised learning, a model is trained to minimize a loss according to manually curated labels, with the most known example being the image-classification task ImageNet [23]. While this approach has merit due to the specificity of the task, and the fact that the manual curations could in fact create a high-quality dataset, its main pitfalls are the cost of manual curation, resulting in a glass ceiling in the sense of dataset size, and the fact that manual labelling is limited by the imagination of the curator. These limitations reduce the models’ abilities to address the world’s complexity and prevent the trained models from generalizing to unseen examples. Interestingly, these same pitfalls were also recently identified as one of the obstacles in classical neuroscience research, which uses hand-picked stimuli, as well as features and labels determined by the experimenters [21].

Addressing this, recent innovations in ANN training have started applying new learning protocols known as “self-supervision” which has delivered great improvements in visual tasks performances [24–29]. One type of such self-supervision is applying image-language multi-modal training, showed great promise in the field of visual perception [24,30] and visual question answering [31–33] with the most publicly familiar model being CLIP [24]. Soon after, preliminary studies in cognition have identified the unique ability to achieve unprecedented performance in predicting fMRI recording in response to visual stimuli [35], effectively crowning image-language models as current state-of-the-art in predicting neural activity, possibly due to the semantic influences of language on the output representations.

In addition, similar training paradigms have been applied to unimodal image-only self-supervised models rivalling CLIP in visual task benchmarks, such as SimCLR and DINOv2 [25,36]. A recent comparison of current state of the art (SOTA) models including CLIP [24], DINO [26], and masked autoencoders [37] displayed a form of representational convergence, showing that increased performance in vision tasks causes the visual representations to converge, regardless of architecture, loss function, and datasets [38]. A first attempt to explain neural recordings using self-supervised models by Konkle et al. [39] showed that some self-supervised models can achieve comparable performance to classification models in predicting fMRI recordings during visual perception tasks. Furthermore, a recent analysis by Prince et al. [40] identified an emergent feature of self-supervised image-only models, which in their broadest definition are trained using a modified image as an input to the model, and a different modified output as an output to the model making the model predict some information evident in the image, having a category selective neurons similar to the category selective regions in the human visual cortex, with said features being highly predictive of the corresponding regions in the human cortex. A recent large-scale study by Conwell et al. [41] directly compared the 3 types of training objectives mentioned in their overall ability to generalize and predict fMRI recordings of the occipital-temporal cortex in a controlled training setting, indicating that any type of self-supervision provides an advantage to predicting neural recordings.

These recent works provide the base for the potential of self-supervision training as a model for the neural mechanisms of visual perception. In this work we attempt to further examine the factors that may underlie the advantage of these self-supervised models over supervised ones. First, based on the importance recent studies assigned to multi-modal self supervised learning, we compared its predictive validity to uni-modal self supervision. Second, given the inherently larger datasets of self supervised relative to supervised learning, we assessed the role of dataset size. Moreover, whereas most previous studies have examined neural prediction of self supervised ANNs with fMRI, here we examined also the neural responses to same stimuli as measured with MEG. This enabled us to uncover the temporal dynamics of ANNs prediction of visual cortex, their similarity to the fMRI data, and their correspondence with the visual cortex hierarchical architecture. To do so we compare current SOTA models trained using either image-language pretraining [24], image-only self-supervision [25,36], and classification trained models [23,42,43] (see Methods) to predict the neural response to natural image stimuli, recorded using both fMRI and MEG collected by Cichy and colleagues [7]. We ask the following questions, is the better prediction of visual-language self-supervised model due to their language supervision or to their self-supervised nature of learning? Does the amount of training data which is typically larger in self-supervised than supervised training account for the superiority of the self-supervised models? At what stage of visual processing do these similarities between self-supervised models and neural responses emerge? To answer these questions, we used models that were all based on vision-transformer (ViT) based architecture, with the same number of trainable parameters, but differed in the nature of training (supervised/self-supervised), multi-modal/unimodal and size of dataset training. By using both fMRI and MEG, we can address these questions from both the spatial and temporal domains.

## Results

### Self-supervised image-language trained models better explain neural representations than supervised image classifiers

To examine whether the representations of the images generated by the self supervised learning (SSL) image-language model, CLIP, explain more variance in the fMRI response to the images than the visual representation generated by a supervised classification model we generated representational dissimilarity matrices (RDMs) based on the algorithm representations across all its layers (see methods) and the fMRI pattern of voxel representations. We then fit a linear model predicting the neural response with either model based on a linear combination of all the layers of the network. The CLIP trained model predicted a significantly greater share of variance in both the neural response in Early Visual Cortex (EVC), as well as in the high-level Ventral Visual Stream (VVS) (*p*_*FDR*_ < 0.05) (see Figure 1A).

**Figure 1:**
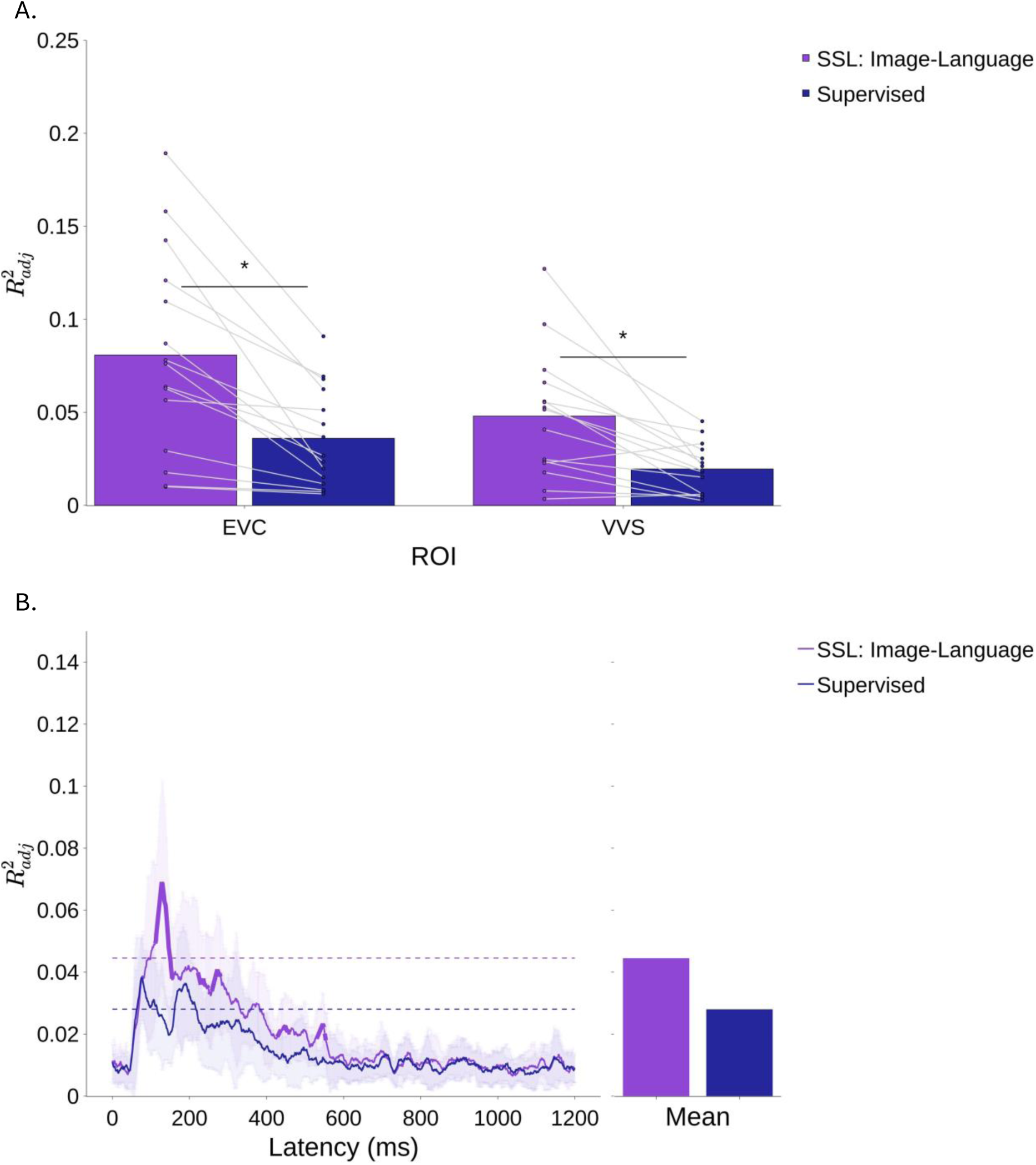
Proportion of variance explained by SSL image-language CLIP (purple) and supervised, ImageNet classification model (blue) of fMRI RDMs (A) and MEG RDMs (B). A. Proportion of explained variance in neural RDMs in early visual cortex (EVC) and ventral visual stream (VVS). Each dot indicates a participant. B. Proportion of explained variance in MEG RDMs. The **bold** section of the line indicates time segments where SSL-vision language model (CLIP) explained a significantly greater proportion of variance than that of the supervised model. Bars and horizontal dashed lines indicate mean 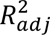 between 80 and 250 ms.

To examine at which time in visual processing CLIP showed this better prediction over ImageNet, we ran a similar analysis on MEG response to the same images for each 1ms. This analysis revealed time segments between 111-158, 222-280 milliseconds where the RDMs of the SSL image-language model were significantly better predictors than the supervised model of the neural RDMs. FDR was used to correct for multiple comparisons (see Figure 1B).

In summary, the SSL image-language model was a better predictor of neural responses of both fMRI and MEG recordings than the supervised DCNN. These findings are in line with recent reports [35]. However, the two models differ in at least three main ways. First, the SSL image-language model is trained on more data. Second, the SSL image-language model is trained in a self-supervised manner. Third, the SSL image-language model is a multi-modal language-vision model, whereas the supervised model is trained only on visual information. We therefore compared the SSL image-language model to a visual-only algorithm that is trained in a self-supervised manner. We then compared multi-modal and uni-modal algorithms trained on a similar size image database.

### Self-supervised image-language and image-only models show complementary advantages over a supervised image classifier

To test if the added value of SSL image-language model (CLIP) is attributed to the semantic influence from language supervision, we examined the proportion of variance explained by a third model, DINOv2, which was trained using only-vision self-supervision on a dataset of similar scale to that of CLIP (142M) (see Methods). DINOv2 explains a significantly larger portion of variance of the fMRI representations than the supervised model, both in the EVC as well as in the VVS (*p*_*FDR*_ < 0.05). While CLIP does not explain a significantly larger portion of variance than DINOv2 in the high level VVS, it does explain a greater proportion of variance in the EVC (*p*_*FDR*_ < 0.05), suggesting that language supervision creates an emergence of visual features that are similar to the EVC. DINOv2 accounted for a significantly larger portion of variance than the supervised model in both EVC and VVS [1,9,10] (See Figure 2).

**Figure 2:**
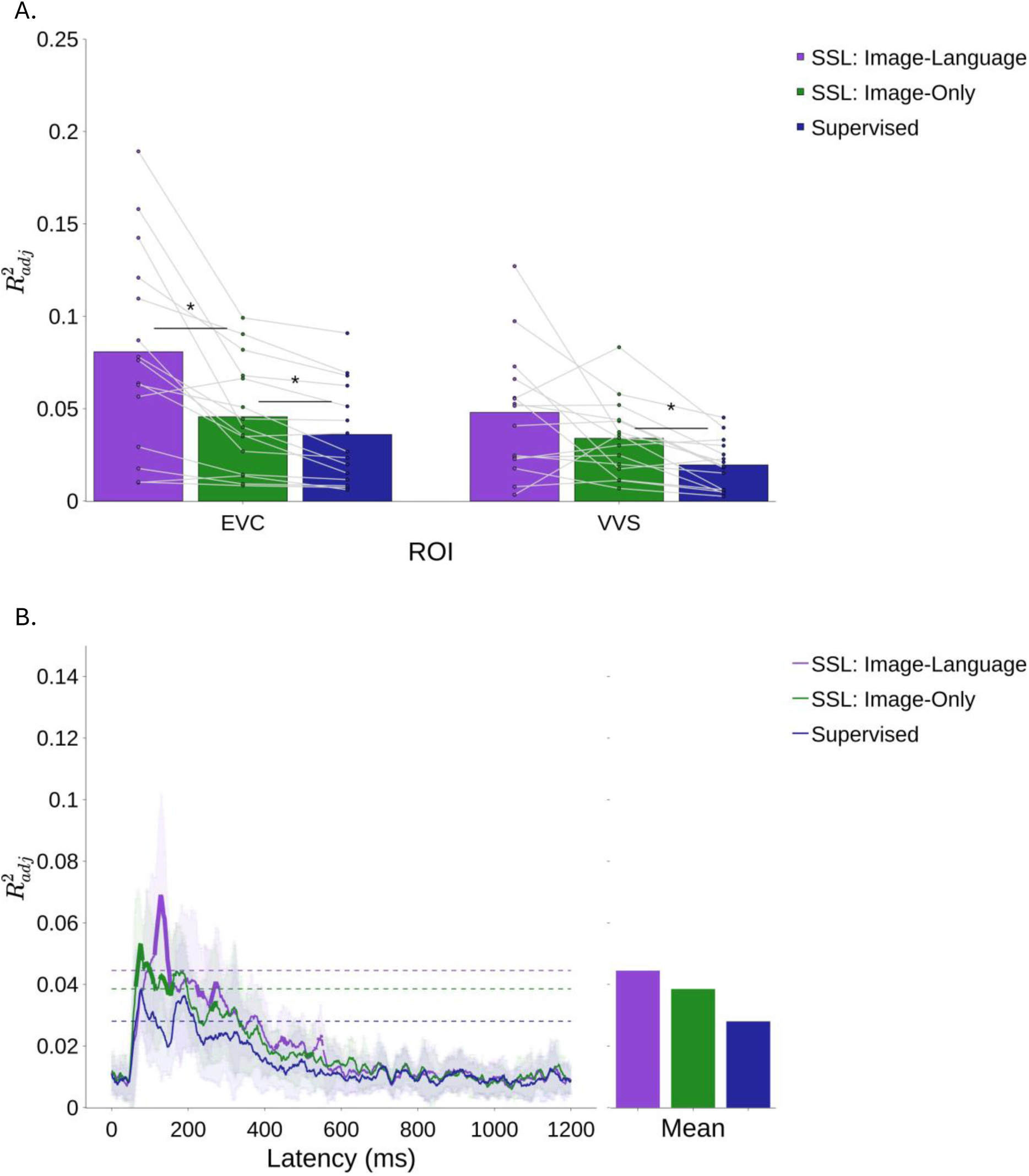
Proportion of variance of fMRI RDMs (A) and MEG RDMs (B) explained by SSL image-language model, CLIP (purple) SSL image-only model, DINOv2 (green) and Supervised classification model (blue). (A) SSL image-only explains significantly more variance than supervised model both at the EVC as well as the VVS, while the SSL image-language model explains more variance than SSL image-only DINOv2 at the EVC. (B) SSL image-only DINOv2 explains significantly more variance than the supervised model at earlier latencies, while CLIP explains greater proportion of variance than the supervised model at later latencies. At no point does SSL image-language model explain a significantly greater proportion of variance than SSL image-only DINOv2 model. Bars and horizontal dashed lines indicate mean 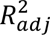 between 80 and 250 ms. * *p*_*FDR*_ < 0.05

A similar analysis on the MEG data showed that SSL image-only DINOv2 explains a significantly larger portion of variance than the supervised model in the time span between 63-81 milliseconds, 83-109 ms, and 126-160 ms, having a large overlap with the timespan explained by SSL image-language, CLIP around 130ms onsets, and prior to that time span. This shows that DINOv2 better explains the neural activity in earlier timespans, while CLIP better explains later onsets.

In summary, when comparing the contributions of DINOv2 and CLIP, we can see that overall, the proportion of variance explained by SSL image-language (CLIP) is not significantly greater than that of SSL image-only (DINOv2), except in fMRI responses in early visual cortex. However, given that CLIP is trained on a much larger dataset (400M) than DiNOv2 (142M), we next examined how dataset size influences predictions of the different models.

### Effects of Dataset size on neural prediction of self-supervised models depend on the type of supervision

To test whether the models trained on larger datasets better predict neural responses, we compared models trained using the same supervision (either SSL image-language, or SSL image-only) trained on datasets of different sizes. First, we compared the OpenAI CLIP variant trained on 400M images [24] used to obtain the previous results, and a CLIP variant trained on 15M image-text pairs from the YFCC dataset [44,45]. CLIP training on a smaller dataset resulted in a model with slightly lesser ability to predict neural activity, as evident in Figure 3. Specifically, the smaller dataset trained model was significantly worse at predicting activity in the EVC in fMRI, whereas the difference in VVS was not statistically significant.

**Figure 3:**
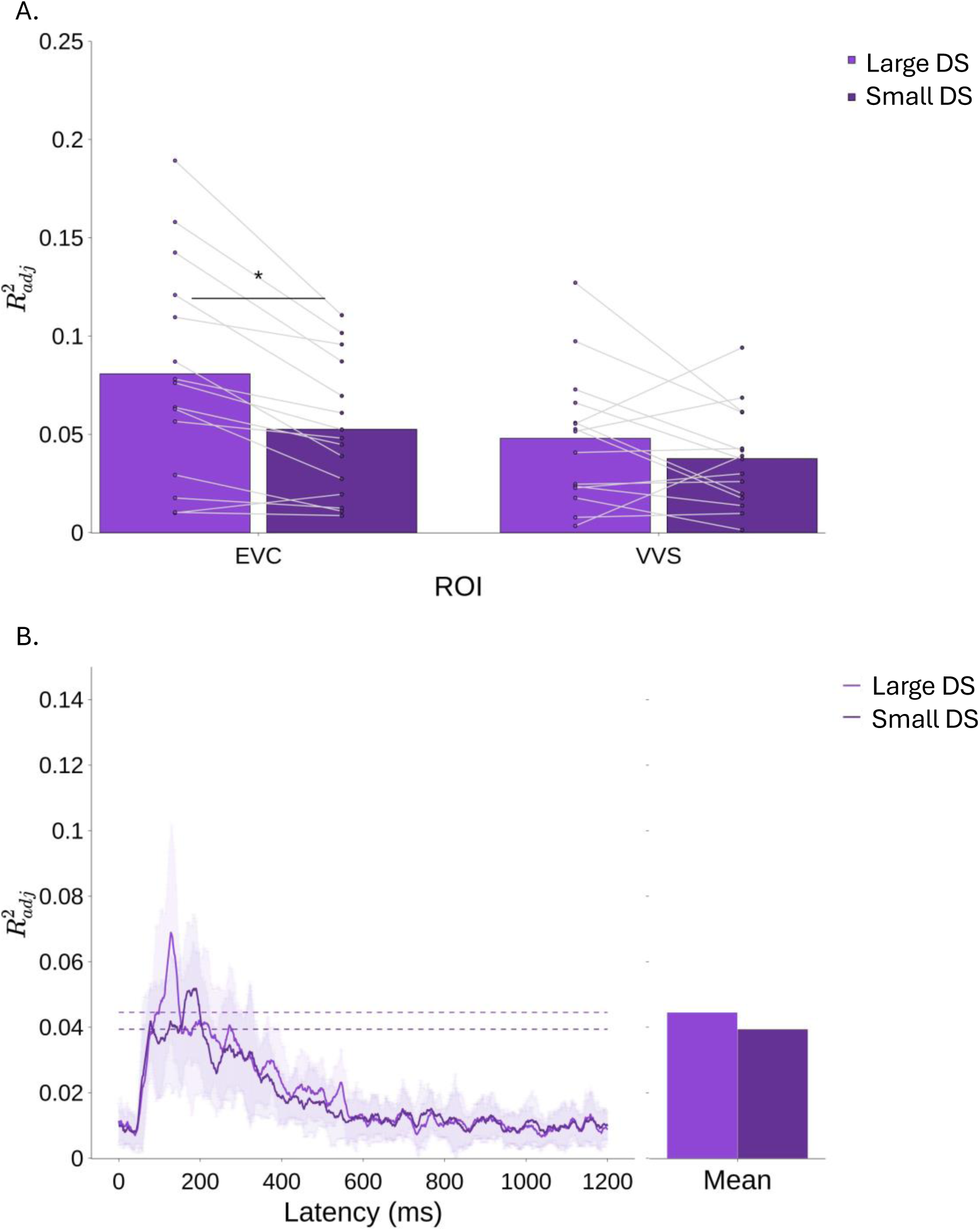
Proportion of variance of fMRI (A) and MEG (B) explained by CLIP trained on large (400M) and small (15M) data sets (DS)). Bars and horizontal dashed lines indicate mean 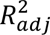 between 80 and 250 ms. * *p*_*FDR*_ < 0.05

Secondly, we compared the ability to predict neural activity using 2 models trained using image-only self-supervision losses, DINOv2, which was trained on 142M images [25], and SimCLR [36] trained on the same 15M images from the YFCC dataset that CLIP was trained on [44,45]. Unlike the case with language supervision, the SimCLR model was significantly better at predicting the fMRI response to images in both the EVC and the VVS, even though it was trained on a much smaller dataset (15M)). Furthermore, the SimCLR model was also significantly better predictive of the MEG responses to the same stimuli set at the interval of 78-124 milliseconds.

**Figure 4:**
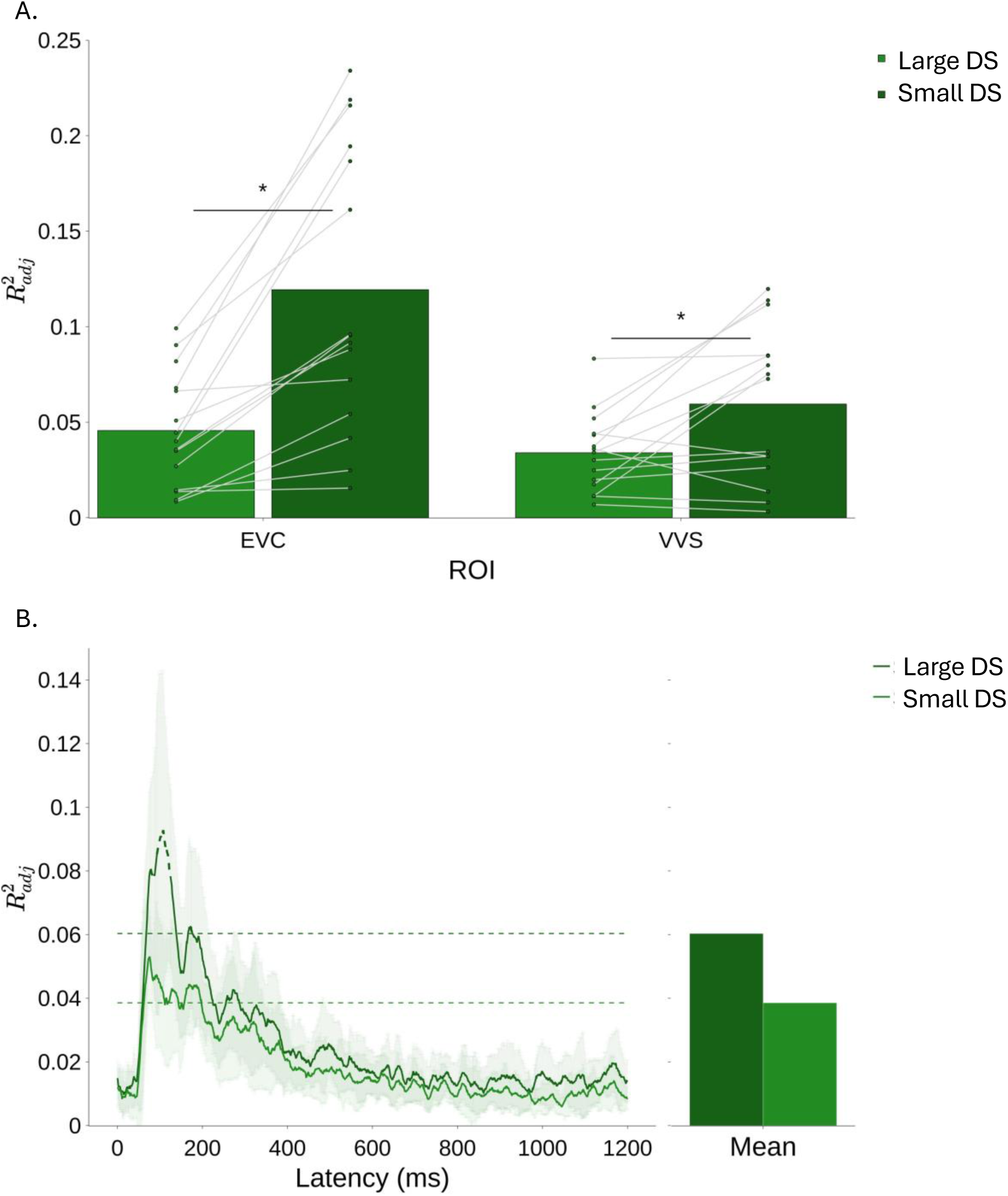
Proportion of variance of fMRI (A) and MEG (B) explained by SSL image only models trained on large data sets (DS) (DINOv2, 142M) and a smaller data set (SimCLR, 15M). Dashed lines in (B) indicate a significantly greater proportion of variance explained by the SimCLR than DINOv2 between 78-124ms. Bars and horizontal dashed lines indicate mean 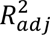 between 80 and 250 ms. * *p*_*FDR*_ < 0.05

It is noteworthy that SimCLR and DinoV2 differ not only in dataset size but also in their loss function (see methods for more details). Still the fact that an SSL model that is trained on substantially smaller dataset can outperform those performed on much larger ones, indicates that increasing dataset size alone cannot guarantee improved correlations with neural data.

Given these effects of dataset size and optimization objective, we examined whether the self-supervised models trained on smaller-dataset variants still possess an advantage at predicting neural activity over the supervised model that was trained on a comparable dataset size. For that we used a DCNN that was trained on the extended dataset ImageNet21k [23] consisting of 21k classes and 15M training images (see Methods).

**Figure 5:**
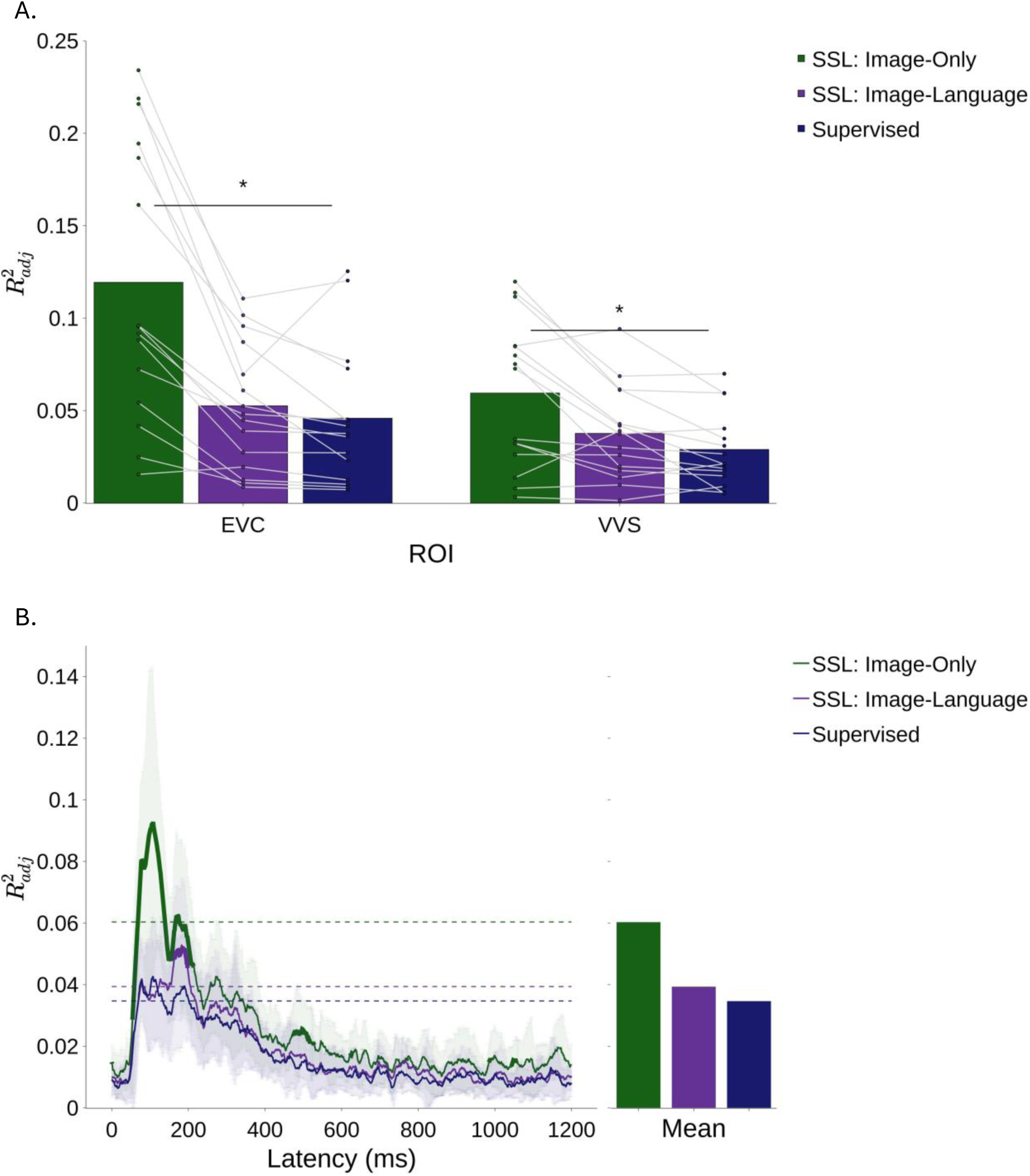
Proportion of variance of fMRI (A) and MEG (B) explained by SSL image-language (CLIP), SSL image only (SimCLR) and supervised classifier, all trained on 15M images. The SSL image only accounted for significant larger proportion of variance than the supervised model in EVC and VVS. SLL image-only model explains a significantly greater proportion of variance than supervised model between 54-214 ms. The SSL image-language model does so at a later offset of 163-198 milliseconds. Bars and horizontal dashed lines indicate mean 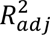 between 80 and 250 ms. * *p*_*FDR*_ < 0.05

Like our previous findings we can see that the self-supervised image-only model (SimCLR) is significantly better than the supervised model trained on the same size of dataset at predicting the EVC as well as the VVS. In contrast with CLIP trained on a large data set (400M), CLIP trained on a smaller data set does not predict a significantly greater portion of variance than the supervised model when predicting fMRI recordings. This is compliant with previous findings showing that a model pretrained using the language supervision on a small dataset display poorer performances in vision tasks [46–48]. Furthermore, we can see that the SSL image-language model is not significantly better than the SSL image only model.

When calculating the proportion of variance explained in the MEG RDMs over time, we can see that SSL image-language model explains a significantly larger proportion of variance than the supervised model at later offsets (163-198 ms), while SSL image only, SimCLR explains a significant proportion of variance between 54-214 ms, a larger timespan that also spans earlier onsets.

### Hierarchical correspondence between visual cortex and self-supervised models

Our analysis so far used a linear combination of all layers of the model to predict brain responses, Next, we wished to break down our analysis to test if different layers explain different stages of processing as recorded by MEG and fMRI. To do so, we examined the proportion of variance explained by the different layers of different self-supervised models. We focus our analysis on CLIP that was trained on 400M images (OpenAI CLIP) and SimCLR that we reported above, due to their better prediction of the neural recordings.

We report the *R*^2^ of each layer of CLIP in figure 6. The earlier layers of CLIP are better at predicting fMRI activity in both the EVC and VVS than later layers. The one unique layer is the last layer of CLIP, trained directly to correspond to embedding of textual description of images, which showed a better fit for the VVS. SimCLR, however best predicts fMRI recordings at the early-intermediate layers. The different layers of both models were better at predicting the EVC than the VVS.

We report the *R*^2^ of each layer of CLIP in figure 7. When predicting MEG recordings, the earlier layers of SimCLR are better at predicting the earlier latencies (50-145ms), while the last layers are better at predicting the later latencies (145-220). The different layers of CLIP do not show the same clear organization, with all layers showing the same competence at predicting the neural activity at all latencies, showing little interaction with the latency, with one exception is the final layer, reaching peak of explained variance at 130ms.

## Discussion

Recent studies have shown the benefit of multi-modal self-supervision models over supervised deep learning models in accounting for human visual processing [35]. The current study aims to assess whether it is the advantages of self-supervision training per se or its multi-modal learning that leads to better prediction of neural responses to images in human visual cortex. We also assessed the contribution of the training dataset size in this process, which is substantially larger in self-supervised than supervised model. Our findings show that unimodal vision self-supervised models are better or as good as multi-modal self-supervised models and that both are better than a supervised model (Figure 2). We found that whereas larger dataset improved prediction for multi-modal models (Figure 3), unimodal self-supervision model trained on smaller dataset were as good or even better in predicting neural activations (Figure 4). Interestingly, a final analysis that examined the contributions of representations of individual layers of the ANNs showed a correspondence between the model hierarchical representations for the multi-modal ANN and the human visual cortex, with the last layer showing specificity in predicting the VVS (Figure 6). While the ability of the different layers of the multi-modal model to predict neural activity through time showed relatively no specificity, with all layers being equally potent at predicting earlier and later onsets of neural response, the image-only self-supervised model showed hierarchical correspondence (Figure 7) with early layers being predictive of earlier onsets, and late layers being predictive of later onsets.

**Figure 6:**
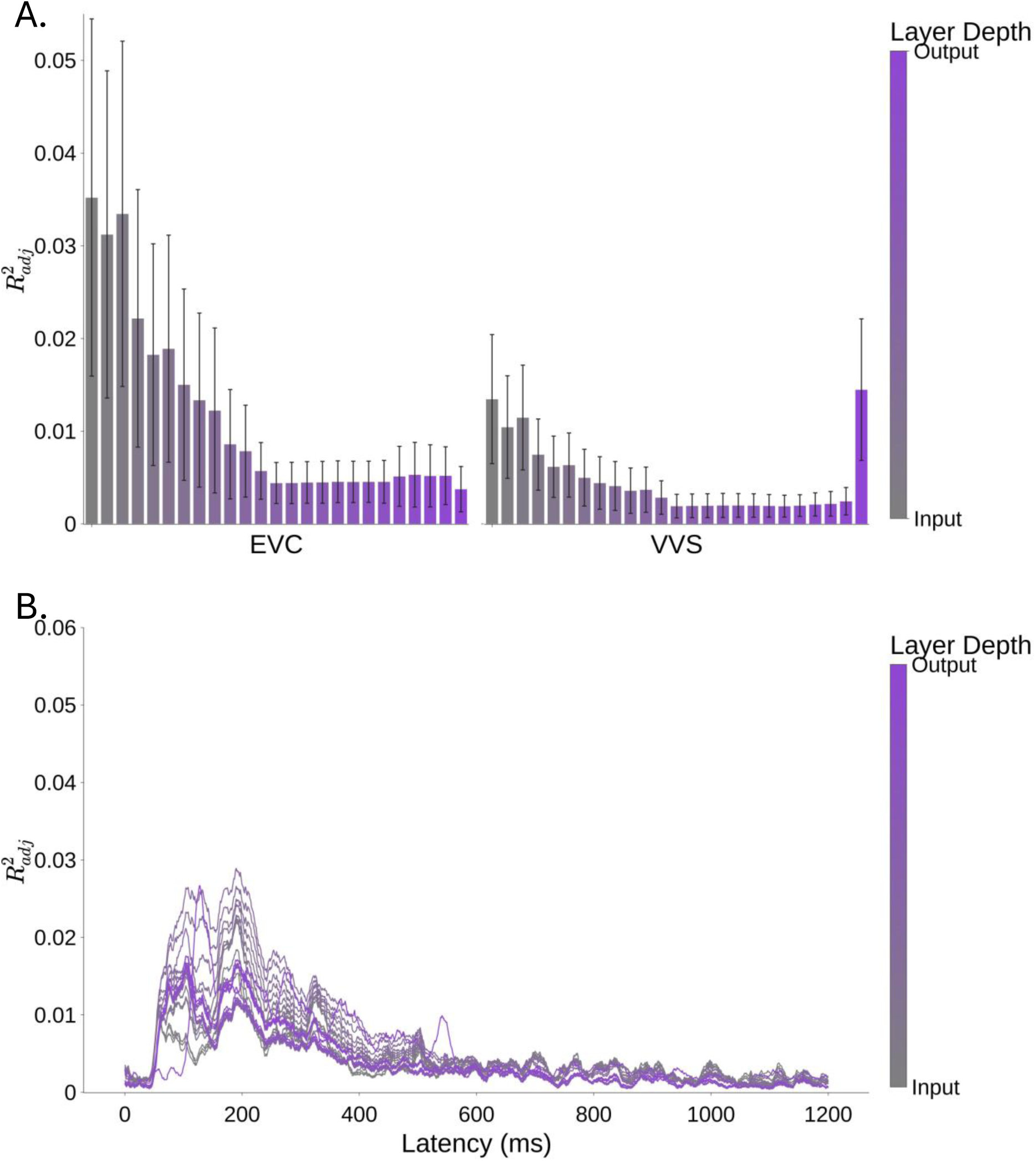
Predictions of individual layers of CLIP (400M) in fMRI (A) and MEG (B). (A) When predicting the EVC and VVS RDMs we can identify an advantage for early CLIP layers over the late layers. Moreover, we observe a sudden surge in proportion of variance that the last layer of CLIP explains in VVS. (B) The different layers of CLIP show increased prediction of neural activity between roughly 100-300ms, with a peak of the final layer around 130ms.

**Figure 7:**
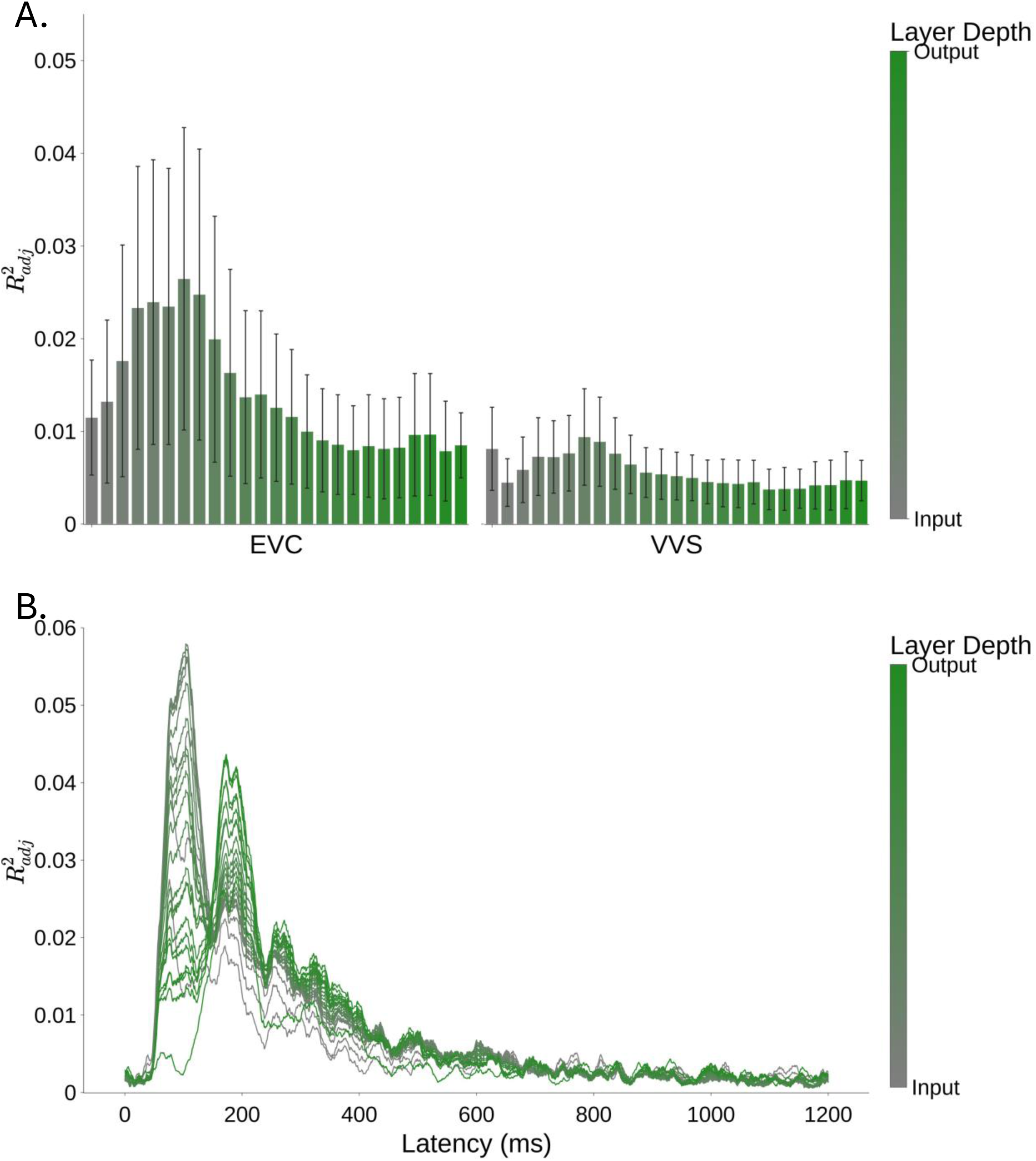
Predictions of individual layers of SimCLR in fMRI (A) and MEG (B). (A) The intermediate-early layers of SimCLR show the best fit of all layers when predicting neural activity in both the EVC and the VVS. (B) Earlier layers of SimCLR are better at predicting earlier latencies (50-145ms) while the later layers of SimCLR are better at predicting later latencies (145-220ms).

### Self-supervision better predicts neural activity than supervised classification

While many previous experiments showed the considerable competence of image classification models in predicting neural activity, we have shown that various types of self-supervision can outperform supervised models in this task. While different training goals showed different advantages over classification training, they were consistent in their ability to outperform supervised learning. Our results show a pattern consistent with previous studies conducted using fMRI recordings [35], and shed new light on temporal dynamics, revealing both image-language supervision, as well as image-only self-supervision, complement classification training, explaining further the MEG recordings at different points in time. This was the case even when the size of the dataset, which is typically larger for self-supervised models, was matched across models (Figure 6).

This conclusion is clearly evident from our analyses, while image-language pretraining explains a significantly larger share of variance than classification training at predicting the neural response to visual stimuli in both MEG and fMRI, it is not unique in its capability, as we have shown that both DINOv2 and SimCLR (image-only self-supervised models) also explain a significantly larger share of variance than classification training in both the EVC and VVS as measured by fMRI, as well as in whole brain measurements in long timespans, as measured by MEG. While CLIP shows significant advantage over image-only training in the EVC when comparing CLIP with DINOv2, this advantage is no longer evident when controlling dataset size and using SimCLR pretraining. Specifically, as detailed below, our findings show that from different factors such as supervision goal and dataset size emerge different similarities to neural recordings.

### Self-supervised models better explain earlier stages of processing

Comparing the SSL models to the findings in Cichy et al (2016) [10], we note that while the early layers of a (convolutional) supervised model reach peak predictivity at latencies as early as 100ms, the visual-only self-supervised model showed a significant advantage at even earlier latencies, reaching a peak at 76ms. Considering the spatio-temporal dynamics identified by [7], this could indicate that the information expressed in the EVC is better extracted from visual-only self-supervision. As previously identified, onsets as early as 80 ms are the earliest to have shown any stimulus-specific neural-response [49]. This to some extent could indicate that learning from visual-only sources (sans language) is a valid candidate for being one of the learning processes of the human brain as previously proposed by [39], as it shows ability to shed light on processing times that have yet to be explained in this scale, while language supervision shows benefit at predicting neural measurements at later onsets. This method could describe neural response in terms of the different emphases placed on different learning objectives at the different stages of perception.

### Training dataset size has different implications on neural predictivity depending on the type of supervision

When trained on a smaller dataset size, the ability of the SSL Image-language (CLIP) to predict neural recordings somewhat diminishes, with a significant drop visible when predicting fMRI recordings at the EVC. This is somewhat expected as previous findings examined the scaling laws of CLIP’s dataset size, identifying that the smaller the number of textual labels, the lesser the performance of the model on various visual tasks [46,50,51]. This is also compliant with the work identifying representational convergence of image models, as they become more competent in common vision tasks [38]. However, this scaling law is not replicated in image-only models, as multiple image-only visual models indeed reach very high performances [36,37,52]. Furthermore, our results show an opposite pattern when comparing DINOv2 and SimCLR, with SimCLR being significantly better at predicting both the EVC and the VVS, as well as in MEG recordings between 78-124ms.

### SSL models’ hierarchies correspond to the brain’s spatial hierarchy

Both learning objectives showed a convergence towards a similar hierarchy as the brain, while the type of optimization seems to govern the type of similarity. Specifically, in both models we can see a decline in explained variance of fMRI as we choose a representation from a deeper layer, with the earlier layers being best predictive of the EVC. However, specifically the last layer of CLIP shows a sudden rise in explained variance of the VVS specifically. As the last layer of CLIP is trained to align with textual descriptions of images, this could indicate a more semantic representation of the VVS in accords with the findings of [20]. When predicting the MEG recordings, a different pattern emerges between the models. Image-Language training did not produce a clear hierarchical correspondence with the MEG recordings, as can be seen in figure 7, all layers were able to explain greater variance between 50 and 350ms. When observing the individual layers of SimCLR we can see a clear hierarchical correspondence, with the early layers being predictive of early latencies, and the last layers being predictive of later latencies. This natural emergent property could indicate a visual-centric learning objective in the human visual cortex.

### Insights into the neural mechanisms of visual perception

One of the main goals of the neuroconnectionist program [21] is to uncover the learning mechanisms that develop cognitive processes. And so, a natural question that arises, is how valid is it to attribute the pattern of results to the training objective? As previously pointed out, a trained model is the sum of various properties, including training objective, optimization algorithm, architecture, dataset and initialization [35]. First, our analysis tackles the dataset diversity by comparing variants trained on a small dataset of 15M images, having comparable dataset size with all models. Second, all models were trained using either Adam [53] or AdamW [54] optimization algorithms as described by [24,42,44], making them relatively comparable. Third, all models are based on the same architecture, Vision Transformer [42]. As noted in [42], a vision transformer does not share the same image-specific inductive-bias as convolutional neural networks, such as translational equivariance. While this in turn makes the architecture somewhat less ‘brain-like’ in the sense of structured convolutions and gradually increasing receptive fields, it could be argued that this property actually makes the attribution of the neural explainability to the training objective as more likely, as it could not be attributed to the architecture’s general aptitude. Furthermore, a comparative analysis of architecture examined the biases of trained vision models showed that transformer-based models had similar biases to human ones as recorded in behavioral experiments [55]. Also, Conwell et al. [41] recently shown that while there is a significant difference in neural predictivity between DCNNs and ViTs, they perform very similarly with the actual difference between them being small. Even so, it is important to address that any difference in architecture could lead to differently converging results, potentially displaying inconsistent results.

Lastly, while this pattern of results identifies different similarities and dissimilarities of the different training mechanisms to the neural visual stream, it is important to reiterate that no single self-supervised model was identified as more similar to the human brain than the others. However, each model shows specific similarities and differences, thus allowing to identify particular components in the learning mechanisms of the human brain.

### Conclusions

In conclusion, whereas DCNNs trained in a supervised manner are the most prevalent ANNs that were used to predict human neural responses in visual cortex, the recent emergences of self-supervised models that enable visual training that is not confined by human labelling offer new ways to understand the type or representations that are generated to visual images in human visual cortex. This includes but is not limited to multi-modal image-language models [35], indicating that unimodal self-supervision also extracts visual information that is aligned with human neural responses. Future studies will further examine the type of features that underlie these similarities between humans and models representations.

## Methods

### Dataset

We used the image dataset from [7] which consisted of 92 natural images containing faces, bodies, hands, places, animals, plants, and inanimate items.

### Neural measurements

We used the publicly available neural data from [7]. To analyse the neural representation at a millisecond resolution we used the precalculated RDMs based on MEG recordings of 16 participants from the first dataset. Each subject was exposed to each image for over 20 times, and the dissimilarity measure was calculated as the Leave-One-Out cross validated linear SVM [56] classification accuracy, classifying which image was displayed based on the MEG measurements at that millisecond.

To calculate the RDM of specific neural ROI, we used fMRI scans from [7]. 15 subjects were exposed to the set 92 images. Scans were directed to record neural responses from the EVC and VVS as explained in [7]. For each stimulus, a searchlight analysis was region. To extract the representation in a specific ROI we extracted all voxels that were defined as part of the ROI according to the probabilistic ROI of [57]. The representational dissimilarity was defined as 1 minus Pearson’s correlation between the t-values of all voxels in the ROI. For further technical details on both neural measurements please refer to [7].

### Models

To predict the neural RDMs we used representations from an ImageNet [23] trained supervised model [42], an image-language self-supervised model trained on 400 million image-language pairs– CLIP [24] and a variant of CLIP trained on a smaller dataset published by [44], and two self-supervised foundational vision model – DINOv2 [25] and SimCLR [36]. While DINOv2 and SimCLR are not trained using the same loss function, both are trained on image-level self supervised, SimCLR using contrastive learning, and DINOv2 using self-distillation without supervision. DINOv2 is further trained using the iBOT supervision [58], performing patch level discrimination. To ensure that one model is not a better fit for predicting the neural RDMs due to implicit biases of architecture, all models were based on the ViT-L architecture [42] Table 1 lists the dataset size, type of training and we cite performance on the ImageNet classification benchmark of the different models from their original publications [24,25,42,44]. Performance on ImageNet classification is stated using 3 different metrics: Without further training the dataset, CLIP models could be used to explicitly examine the similarities between images and class textual representations, allowing for 0-shot classification. As DINOv2 is an SSL trained model that does not explicitly support this, we report K-nearest-neighbours’ performance. A second metric is linear probing, where the model is used as a frozen image encoder and only a linear classifier is trained to measure the model’s classification capabilities. Third, we report performance after finetuning the entire model on ImageNet data.

**Table 1:**
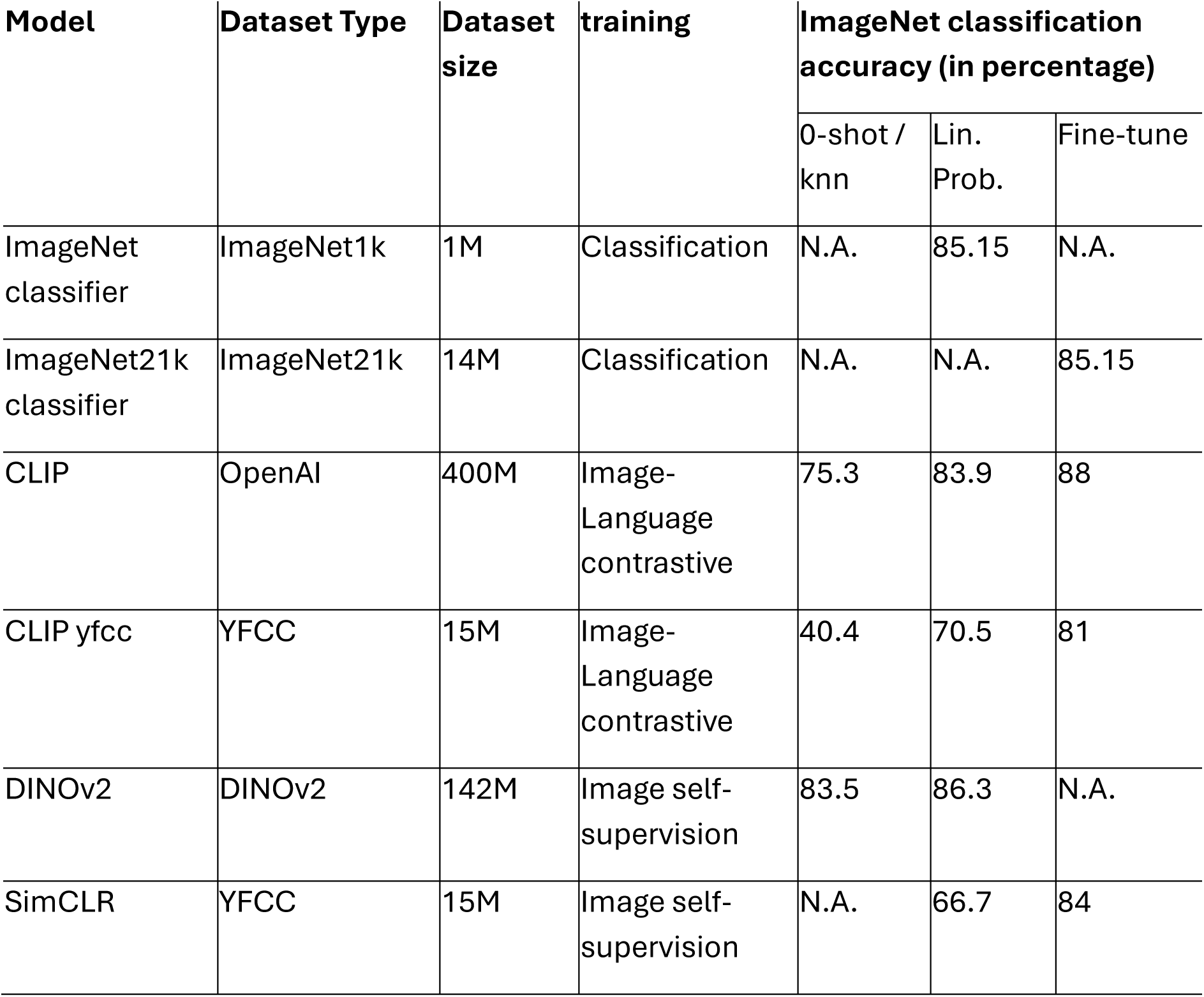
models used in the experiments. Includes the dataset name, size, type of training, and ImageNet classification accuracy, measured using 0-shot \ k nearest neighbors, linear probing, or full finetuning.

| Model | Dataset Type | Dataset size | training | ImageNet classification accuracy (in percentage) |  |  |
| --- | --- | --- | --- | --- | --- | --- |
|  |  |  |  | 0-shot / knn | Lin. Prob. | Fine-tune |
| ImageNet classifier | ImageNet1k | 1M | Classification | N.A. | 85.15 | N.A. |
| ImageNet21k classifier | ImageNet21k | 14M | Classification | N.A. | N.A. | 85.15 |
| CLIP | OpenAI | 400M | Image-Language contrastive | 75.3 | 83.9 | 88 |
| CLIP yfcc | YFCC | 15M | Image-Language contrastive | 40.4 | 70.5 | 81 |
| DINOv2 | DINOv2 | 142M | Image self-supervision | 83.5 | 86.3 | N.A. |
| SimCLR | YFCC | 15M | Image self-supervision | N.A. | 66.7 | 84 |

### Model RDM calculation

As the same information could emerge from different models at different stages of processing, we extracted the representation after every transformer-encoder block in addition to the output representations from each model. To calculate the dissimilarities between the representations at layer ℓ of each pair of images, we calculated the cosine dissimilarity between their embeddings.

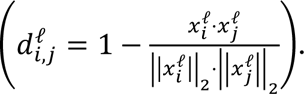

### Multi-layer RSA analysis

To measure how well the dissimilarity ratings of a model can predict the neural dissimilarity ratings, for each subject we fitted a linear regression model predicting the dissimilarity ratings using the set of dissimilarity ratings from all layers and measured the explained portion of variance (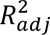). Specifically, to predict the MEG dissimilarity ratings at time *τ* of subject *i* we fitted the linear model minimizing 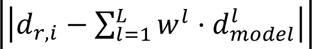. To predict the fMRI dissimilarity ratings at ROI *r* of subject *i* we fitted the linear model minimizing 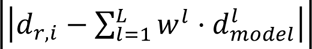.

As the number of layers in ViT-L is relatively large (24) the number of features used to predict the dissimilarity ratings is very large as well, which could easily lead to overfitting. To counter this effect, we used PCA to reduce the number of features for every model.

For the PCA regularization, we first observed the percentage of variance explained by each component, choosing the number of components such that the transformed features will contain at least 95% of explained variance for every model, resulting in using 6 features that were used for predicting the neural RDMs (see supp Figure 1).

### Significance testing

To examine whether a model is significantly better at predicting the neural measurements than all other models, we used the Wilcoxon signed-rank test [59] for paired samples, comparing the percentage of explained variance of each subject at every millisecond \ ROI is greater for one model in comparison to another.

We then applied FDR correction [60] on all calculated p values to correct for multiple comparisons and consider a significant preference only when *p*_*FDR*_ < 0.05.

### Model prediction estimation

To measure the ability of emergent features to explain neural activity, we extracted the representations from every intermediate layer of the model, as well as the last layer. We then calculated the dissimilarity ratings of the embeddings of the input images at every layer. As consecutive layers have very correlated representations, we performed PCA on the similarity ratings of a model and chose the first 6 principal components for analysis. We then fit a linear regression model to predict the neural dissimilarity of a particular subject at a particular ROI for dissimilarity ratings based on fMRI measurements, or at a particular millisecond for dissimilarity ratings based on MEG measurements. To test if one model is better predictive of the neural measurements than another model at a particular time or ROI, we performed a repeated measurements Wilcoxon rank-permutation test [61], comparing the proportion of variance explained by one model for a specific subject, with the proportion of variance explained by another model for the same subject. To correct for multiple comparisons, we performed a false-discovery-rate correction [60] on the calculated p-values.

## References

1. Hubel DH, Wiesel TN. Receptive fields, binocular interaction and functional architecture in the cat’s visual cortex. J Physiol. 1962;160: 106. doi:10.1113/JPHYSIOL.1962.SP006837

2. Hegdé J, Van Essen DC. Selectivity for Complex Shapes in Primate Visual Area V2. The Journal of Neuroscience. 2000;20. Available:www.jneurosci.org. http://www.jneurosci.org/cgi/content/full/3976

3. Wilson HR, Wilkinson F. From orientations to objects: Configural processing in the ventral stream. J Vis. 2015;15: 4–4. doi:10.1167/15.7.4

4. Gross CG, Rocha-Miranda CE, Bender DB. Visual properties of neurons in inferotemporal cortex of the Macaque. J Neurophysiol. 1972;35: 96–111. doi:10.1152/JN.1972.35.1.96

5. Hung CP, Kreiman G, Poggio T, DiCarlo JJ. Fast readout of object identity from macaque inferior temporal cortex. Science (1979). 2005;310: 863–866. doi:10.1126/SCIENCE.1117593/SUPPL_FILE/HUNG.SOM.PDF

6. Grill-Spector K, Malach R. The human visual cortex. Annu Rev Neurosci. 2004;27: 649–677. doi:10.1146/ANNUREV.NEURO.27.070203.144220

7. Cichy RM, Pantazis D, Oliva A. Similarity-Based Fusion of MEG and fMRI Reveals Spatio-Temporal Dynamics in Human Cortex During Visual Object Recognition. Cereb Cortex. 2016;26: 3563–3579. doi:10.1093/cercor/bhw135

8. Cichy RM, Pantazis D, Oliva A. Resolving human object recognition in space and time. Nature Neuroscience 2014 17:3. 2014;17: 455–462. doi:10.1038/nn.3635

9. Khaligh-Razavi SM, Kriegeskorte N. Deep Supervised, but Not Unsupervised, Models May Explain IT Cortical Representation. PLoS Comput Biol. 2014;10: e1003915. doi:10.1371/JOURNAL.PCBI.1003915

10. Cichy RM, Khosla A, Pantazis D, Torralba A, Oliva A. Comparison of deep neural networks to spatio-temporal cortical dynamics of human visual object recognition reveals hierarchical correspondence. Scientific Reports 2016 6:1. 2016;6: 1–13. doi:10.1038/srep27755

11. Layton OW, Steinmetz ST. Convolutional neural networks as a model for brain area MSTd. bioRxiv. 2024; 2024.01.26.577393. doi:10.1101/2024.01.26.577393

12. Soulos P, Isik L. Disentangled deep generative models reveal coding principles of the human face processing network. bioRxiv. 2023; 2023.02.15.528489. doi:10.1101/2023.02.15.528489

13. Tuckute G, Feather J, Boebinger D, McDermott JH. Many but not all deep neural network audio models capture brain responses and exhibit correspondence between model stages and brain regions. PLoS Biol. 2023;21: e3002366. doi:10.1371/JOURNAL.PBIO.3002366

14. Cichy RM, Kaiser D. Deep Neural Networks as Scientific Models. Trends Cogn Sci. 2019;23: 305–317. doi:10.1016/J.TICS.2019.01.009/ASSET/11EA1E97-58C6-46A2-A2A2-147136637450/MAIN.ASSETS/GR2.JPG

15. Richards BA, Lillicrap TP, Beaudoin P, Bengio Y, Bogacz R, Christensen A, et al. A deep learning framework for neuroscience. Nature Neuroscience 2019 22:11. 2019;22: 1761–1770. doi:10.1038/s41593-019-0520-2

16. Storrs KR, Kietzmann TC, Walther A, Mehrer J, Kriegeskorte N. Diverse Deep Neural Networks All Predict Human Inferior Temporal Cortex Well, After Training and Fitting. J Cogn Neurosci. 2021;33: 2044–2064. doi:10.1162/JOCN_A_01755

17. Ratan Murty NA, Bashivan P, Abate A, DiCarlo JJ, Kanwisher N. Computational models of category-selective brain regions enable high-throughput tests of selectivity. Nature Communications 2021 12:1. 2021;12: 1–14. doi:10.1038/s41467-021-25409-6

18. Dwivedi K, Bonner MF, Cichy RM, Roig G. Unveiling functions of the visual cortex using task-specific deep neural networks. PLoS Comput Biol. 2021;17: e1009267. doi:10.1371/JOURNAL.PCBI.1009267

19. Jozwik KM, Kietzmann TC, Cichy RM, Kriegeskorte N, Mur M. Deep Neural Networks and Visuo-Semantic Models Explain Complementary Components of Human Ventral-Stream Representational Dynamics. Journal of Neuroscience. 2023;43: 1731–1741. doi:10.1523/JNEUROSCI.1424-22.2022

20. Devereux BJ, Clarke A, Tyler LK. Integrated deep visual and semantic attractor neural networks predict fMRI pattern-information along the ventral object processing pathway. Scientific Reports 2018 8:1. 2018;8: 1–12. doi:10.1038/s41598-018-28865-1

21. Doerig A, Sommers RP, Seeliger K, Richards B, Ismael J, Lindsay GW, et al. The neuroconnectionist research programme. Nature Reviews Neuroscience 2023 24:7. 2023;24: 431–450. doi:10.1038/s41583-023-00705-w

22. Simony E, Grossman S, Malach R. Brain-machine convergent evolution: Why finding parallels between brain and artificial systems is informative. Proc Natl Acad Sci U S A. 2024;121: e2319709121. doi:10.1073/PNAS.2319709121/ASSET/2C05C90C-BB06-47D9-8ABE-401A97CEB85A/ASSETS/IMAGES/LARGE/PNAS.2319709121FIG04.JPG

23. Deng J, Dong W, Socher R, Li L-J, Kai Li, Li Fei-Fei. ImageNet: A large-scale hierarchical image database. 2009 IEEE Conference on Computer Vision and Pattern Recognition. IEEE; 2009. pp. 248–255. doi:10.1109/CVPR.2009.5206848

24. Radford A, Kim JW, Hallacy C, Ramesh A, Goh G, Agarwal S, et al. Learning Transferable Visual Models From Natural Language Supervision. Proc Mach Learn Res. 2021;139: 8748–8763. Available: https://arxiv.org/abs/2103.00020v1

25. Oquab M, Darcet T, Moutakanni T, Vo H V, Szafraniec M, Khalidov V, et al. DINOv2: Learning Robust Visual Features without Supervision. 2023 [cited 14 Feb 2024]. Available: https://arxiv.org/abs/2304.07193v2

26. Caron M, Touvron H, Misra I, Jégou H, Mairal J, Bojanowski P, et al. Emerging Properties in Self-Supervised Vision Transformers. 2021 [cited 6 Feb 2024]. Available: http://arxiv.org/abs/2104.14294

27. Feichtenhofer C, Fan H, Li Y, He K, Ai M. Masked Autoencoders As Spatiotemporal Learners. Adv Neural Inf Process Syst. 2022;35: 35946–35958. Available: https://github.com/facebookresearch/mae_st

28. Huang P-Y, Sharma V, Xu H, Ryali C, Fan H, Li Y, et al. MAViL: Masked Audio-Video Learners. 2023. Available: https://github.com/facebookresearch/MAViL.

29. He K, Chen X, Xie S, Li Y, Dollar P, Girshick R. Masked Autoencoders Are Scalable Vision Learners. Proceedings of the IEEE Computer Society Conference on Computer Vision and Pattern Recognition. 2021;2022-June: 15979–15988. doi:10.1109/CVPR52688.2022.01553

30. Cherti M, Beaumont R, Wightman R, Wortsman M, Ilharco G, Gordon C, et al. Reproducible scaling laws for contrastive language-image learning. 2022; 2818– 2829. doi:10.1109/cvpr52729.2023.00276

31. Li J, Li D, Savarese S, Hoi S. BLIP-2: Bootstrapping Language-Image Pre-training with Frozen Image Encoders and Large Language Models. Proc Mach Learn Res. 2023;202: 20351–20383. Available: https://arxiv.org/abs/2301.12597v3

32. Li J, Li D, Xiong C, Hoi S. BLIP: Bootstrapping Language-Image Pre-training for Unified Vision-Language Understanding and Generation. Proc Mach Learn Res. 2022;162: 12888–12900. Available: https://arxiv.org/abs/2201.12086v2

33. Mokady R, Hertz A, Bermano AH. ClipCap: CLIP Prefix for Image Captioning. 2021 [cited 16 Jan 2024]. Available: http://arxiv.org/abs/2111.09734

34. Shoham A, Grosbard I, Ger Y, Kossovsky S, Barnahor T, Yovel G. The representational geometry of images and concepts in perception and memory. In: Vision Sciences Society Conference [Internet]. 17 May 2022 [cited 1 Sep 2022]. Available: https://www.visionsciences.org/presentation/?id=3889

35. Wang AY, Kay K, Naselaris T, Tarr MJ, Wehbe L. Better models of human high-level visual cortex emerge from natural language supervision with a large and diverse dataset. Nature Machine Intelligence 2023 5:12. 2023;5: 1415–1426. doi:10.1038/s42256-023-00753-y

36. Chen T, Kornblith S, Norouzi M, Hinton G. A Simple Framework for Contrastive Learning of Visual Representations. 37th International Conference on Machine Learning, ICML 2020. 2020;PartF168147-3: 1575–1585. Available: https://arxiv.org/abs/2002.05709v3

37. He K, Chen X, Xie S, Li Y, Dollár P, Girshick R. Masked Autoencoders Are Scalable Vision Learners. 2022. pp. 16000–16009.

38. Huh M, Cheung B, Wang T, Isola P. The Platonic Representation Hypothesis. 2024 [cited 20 Sep 2024]. Available: https://arxiv.org/abs/2405.07987v5

39. Konkle T, Alvarez GA. A self-supervised domain-general learning framework for human ventral stream representation. Nature Communications 2022 13:1. 2022;13: 1–12. doi:10.1038/s41467-022-28091-4

40. Prince JS, Alvarez GA, Konkle T. Contrastive learning explains the emergence and function of visual category-selective regions. Sci Adv. 2024;10: 1776. doi:10.1126/SCIADV.ADL1776

41. Conwell C, Prince JS, Kay KN, Alvarez GA, Konkle T. A large-scale examination of inductive biases shaping high-level visual representation in brains and machines. Nature Communications 2024 15:1. 2024;15: 1–18. doi:10.1038/s41467-024-53147-y

42. Dosovitskiy A, Beyer L, Kolesnikov A, Weissenborn D, Zhai X, Unterthiner T, et al. An Image is Worth 16×16 Words: Transformers for Image Recognition at Scale. ICLR 2021 - 9th International Conference on Learning Representations. 2020 [cited 7 Aug 2023]. Available: https://arxiv.org/abs/2010.11929v2

43. Russakovsky O, Deng J, Su H, Krause J, Satheesh S, Ma S, et al. ImageNet Large Scale Visual Recognition Challenge. Int J Comput Vis. 2014;115: 211–252. doi:10.1007/s11263-015-0816-y

44. Mu N, Kirillov A, Wagner D, Xie S. SLIP: Self-supervision meets Language-Image Pre-training. Lecture Notes in Computer Science (including subseries Lecture Notes in Artificial Intelligence and Lecture Notes in Bioinformatics). 2021;13686 LNCS: 529–544. doi:10.1007/978-3-031-19809-0_30

45. Thomee B, Shamma DA, Friedland G, Elizalde B, Ni K, Poland D, et al. YFCC100M: The New Data in Multimedia Research. Commun ACM. 2015;59: 64–73. doi:10.1145/2812802

46. Yang C, An Z, Huang L, Bi J, Yu X, Yang H, et al. CLIP-KD: An Empirical Study of CLIP Model Distillation. 2023 [cited 4 Sep 2024]. Available: https://arxiv.org/abs/2307.12732v2

47. Wu K, Peng H, Zhou Z, Xiao B, Liu M, Yuan L, et al. TinyCLIP: CLIP Distillation via Affinity Mimicking and Weight Inheritance.

48. Li Z, Deepmind G, Xie C, Dogus Cubuk E. Scaling (Down) CLIP: A Comprehensive Analysis of Data, Architecture, and Training Strategies.

49. Carlson T, Tovar DA, Alink A, Kriegeskorte N. Representational dynamics of object vision: The first 1000 ms. J Vis. 2013;13: 1–1. doi:10.1167/13.10.1

50. Li Z, Deepmind G, Xie C, Dogus Cubuk E. Scaling (Down) CLIP: A Comprehensive Analysis of Data, Architecture, and Training Strategies. 2024 [cited 29 Sep 2024]. Available: https://arxiv.org/abs/2404.08197v2

51. Wu K, Peng H, Zhou Z, Xiao B, Liu M, Yuan L, et al. TinyCLIP: CLIP Distillation via Affinity Mimicking and Weight Inheritance. Proceedings of the IEEE International Conference on Computer Vision. 2023; 21913–21923. doi:10.1109/ICCV51070.2023.02008

52. Chen M, Radford A, Child R, Wu J, Jun H, Luan D, et al. Generative Pretraining from Pixels. Proceedings of the 37 th International Conference on Machine Learning. 2020 [cited 30 Sep 2024]. Available: https://cdn.openai.com/papers/Generative_Pretraining_from_Pixels_V1_ICML.pdf

53. Kingma DP, Ba JL. Adam: A method for stochastic optimization. 3rd International Conference on Learning Representations, ICLR 2015 - Conference Track Proceedings. 2015.

54. Loshchilov I, Hutter F. Decoupled Weight Decay Regularization. 7th International Conference on Learning Representations, ICLR 2019. 2017 [cited 1 Oct 2024]. Available: https://arxiv.org/abs/1711.05101v3

55. Tuli S, Dasgupta I, Grant E, Griffiths TL. Are Convolutional Neural Networks or Transformers more like human vision? Proceedings of the 43rd Annual Meeting of the Cognitive Science Society: Comparative Cognition: Animal Minds, CogSci 2021. 2021; 1844–1850. Available: https://arxiv.org/abs/2105.07197v2

56. Cortes C, Vapnik V, Saitta L. Support-vector networks. Machine Learning 1995 20:3. 1995;20: 273–297. doi:10.1007/BF00994018

57. Wang L, Mruczek REB, Arcaro MJ, Kastner S. Probabilistic Maps of Visual Topography in Human Cortex. Cerebral Cortex. 2015;25: 3911–3931. doi:10.1093/CERCOR/BHU277

58. Zhou J, Wei C, Wang H, Shen W, Xie C, Yuille A, et al. iBOT: Image BERT Pre-Training with Online Tokenizer. ICLR 2022 - 10th International Conference on Learning Representations. 2021 [cited 15 Feb 2024]. Available: https://arxiv.org/abs/2111.07832v3

59. Wilcoxon F. Individual Comparisons by Ranking Methods. Biometrics Bulletin. 1945;1.

60. Benjamini Y, Yekutieli D. The control of the false discovery rate in multiple testing under dependency. 101214/aos/1013699998. 2001;29: 1165–1188. doi:10.1214/AOS/1013699998

61. Wilcoxon F. Individual Comparisons by Ranking Methods. Biometrics Bulletin. 1945;1: 80. doi:10.2307/3001968

