## Supplemental information for "Self-supervision deep learning models are better models of human high-level visual cortex: The roles of multi-modality and dataset training size"

Supplementary

PCA analysis of each model. As described in the methods section, to reduce the number of predictors and reduce overfit we performed PCA on the dissimilarity ratings of all layers of a particular model, and chose the first 6 components, explaining at least 99.7% of variance for each model.

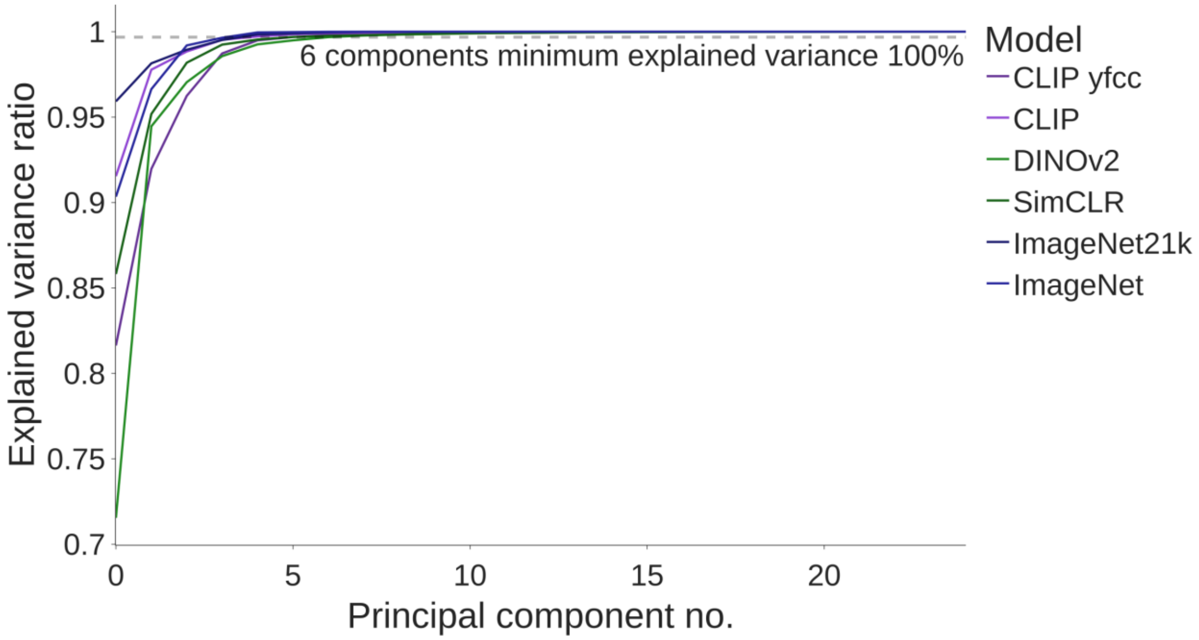


Figure 1: Proportion of variance explained (y axis) by each component (x axis) of each model (trend color).
